# Towards Sparse Causal Features for Zero-shot Mutation Effect Prediction in a Protein Language Model

**DOI:** 10.64898/2026.08.28.747907

**Authors:** Saishradha Mohanty, Manya Phutela, Anna G. Green

## Abstract

Protein language models (pLMs) such as ESM-2 achieve strong zero-shot mutation-effect prediction, yet the internal computations supporting these predictions remain poorly understood. We introduce a sparse feature circuit framework that combines sparse autoencoders, integrated-gradients attribution, and activation patching to identify the latent features that causally mediate zero-shot mutation effect prediction in ESM-2 650M. We evaluate this framework over 67 mutations ranging from strongly deleterious to weakly deleterious in the DNAJA1 J-domain, where ESM-2 predictions agree strongly with deep mutational scanning measurements. We find that circuits selected by indirect effect recover the model’s predictions more efficiently and provide more informative biological explanations than those selected by raw activation changes, showing that activation magnitude does not necessarily reflect causal importance. We find that related substitutions reuse substantial portions of their recovered circuits, ranging from 40% to 75%, and that the shared features often represent residues in three-dimensional contact with the mutation site. To our knowledge, our work provides the first causal, feature-level account of zero-shot mutation effect prediction in a pLM.

## 1 Introduction

Predicting the effects of protein mutations is a central problem in computational biology, with direct applications in protein engineering and disease variant interpretation. Supervised methods can predict the effects of amino acid substitutions on protein function, but require experimental or clinical training data in order to be trained. Zero-shot mutation-effect prediction provides a useful alternative for predicting the effects of mutations in proteins for which such data is unavailable. By learning a probability distribution over sequence space, models can assign mutation effect scores without mutation and assay-specific supervision. Protein language models, which are trained on hundreds of millions of natural sequences, have achieved strong performance on variant-effect prediction benchmarks (Meier et al. 2021; Notin et al. 2023). ESM-2 650M (Lin et al. 2023) in particular achieves strong rank correlation with experimentally measured effects across many proteins and assays. Yet, a high zero-shot score does not explain what features a model has learned, whether those features are biologically meaningful, or how a model computes its prediction from those features.

Recent mechanistic interpretability work has begun to characterize *what* features pLMs represent. Sparse autoencoders (SAEs) decompose dense residual-stream activations into a larger dictionary of sparsely activating, approximately monosemantic features (Bricken et al. 2023; Cunningham et al. 2024). Application of SAEs to pLMs has recovered features that activate on protein domains, structural motifs, binding sites, and family-specific signals (Simon and Zou 2025; Gujral et al. 2025; Adams et al. 2025; Villegas Garcia and Ansuini 2025; Liu et al. 2026). Agentic systems have been developed to automate the process of matching the SAE latents to biological annotations (Candido et al. 2026; Gao et al. 2025b; Pearce et al. 2026). However, these analyses reveal how a model input influences the activation of internal interpretable features, but not whether and how those features are *recruited* when the model makes a specific prediction.

Methods from causal analysis allow attribution of network behavior to specific model components. Circuit discovery is a type of causal analysis that seeks to discover a minimal set of internal model representations and computations that determine a model’s behavior (Wang et al. 2023; Marks et al. 2025). Circuit discovery has been applied to supervised mutation effect prediction (Tsui et al. 2026) and contact prediction (Nainani et al. 2025), identifying sparse features with causal contributions, but has not yet been applied to unsupervised mutation effect prediction. ESMZero (Wang et al. 2026) steers SAE features to improve zero-shot mutation effect prediction, but they identify candidate features based on their activation changes between wild-type and mutant sequences, not their causal contributions.

### Contributions

Here, we provide the first sparse feature-circuit analysis of zero-shot mutation effect prediction in a protein language model. Using causal interventions on SAE latents, we identify the features recruited for individual predictions and assess whether they encode biologically plausible explanations. We use single substitutions in DNAJA1 as computed by ESM-2 650M as our case study.

- We introduce a sparse feature-circuit discovery framework combining indirect-effect attribution with Jensen–Shannon recovery to identify causally relevant token–latent features.
- Across 67 substitutions in four DNAJA1 residue positions, we show that indirect-effect ranking recovers the model computation more reliably and efficiently than activation-change ranking.
- We evaluate a customized version of ToolUniverse’s variant-interpretation tools and find that corpus-derived descriptions are limited in their ability to explain protein-specific mutation effects.
- We provide an in-depth analysis of the recovered circuits, showing that related substitutions reuse sparse features and that shared circuits at buried sites are preferentially enriched in three-dimensional contacts around the mutation site.

## 2 Related Work

### Protein language models for mutation effect prediction

Transformer-based protein language models trained on large sequence databases via masked language modeling learn representations that encode evolutionary, structural, and functional information (Rives et al. 2021; Lin et al. 2023). Meier et al. showed that masked-marginal log-likelihood ratios enable accurate zero-shot mutation effect prediction across diverse proteins and assays without task-specific supervision (Meier et al. 2021). ProteinGym standardized evaluation across hundreds of deep mutational scanning datasets and demonstrated strong performance for ESM-family models (Notin et al. 2023). Yet, zero-shot scores do not reveal the internal reasoning behind predictions: *why does the model assign a particular substitution a deleterious score?*

### Interpretability of protein language models

Early mechanistic analyses relied on attention maps and probing classifiers, revealing that pLM attention heads partially align with residue–residue contacts, secondary structure, and functional annotations (Vig et al. 2021; Rao et al. 2021). However, correlating attention weights with biological labels does not identify which model components causally drive a specific output—a limitation well established in the NLP interpretability literature (Jain and Wallace 2019; Wiegreffe and Pinter 2019). More recently, sparse autoencoders (SAEs) have been used to decompose pLM residual-stream activations into a larger, more interpretable feature dictionary. Building on earlier work showing that neural networks represent more features than neurons via superposition (Elhage et al. 2022), SAE-based approaches (Bricken et al. 2023; Cunningham et al. 2024; Gao et al. 2025a) recover approximately monosemantic features that can be annotated with human-interpretable concepts. Adams et al. trained TopK SAEs on multiple residual-stream layers of ESM-2 and showed that many latents activate on specific motifs, domains, and structural patterns (Adams et al. 2025); InterPLM demonstrates that ESM-2 SAE features correspond to biological concepts including protein domains, structural annotations, and functional sites (Simon and Zou 2025); and (Gujral et al. 2025) provide additional evidence for the utility of SAE representations. Critically, all these studies characterize what features a pLM *represents* across its full training distribution—none identifies which features are *recruited* for a specific downstream computation such as mutation-effect prediction.

### Interpreting protein language models using agents

Recently, agentic systems have been used to automate the interpretation of zero-shot mutation-effect predictions from biological foundation models. ToolUniverse’s (Gao et al. 2025b) `ESM_explain_variant_mechanism` compares wildtype and mutant ESM-C (Candido et al. 2026) SAE activations and describes affected features using annotations inferred from a multi-protein reference panel. Similarly, EVEE probes Evo 2 embeddings for biological annotations, quantifies variant-induced changes in these signals, and using a Claude reasoning model synthesizes them with variant and gene context into mechanistic hypotheses (Pearce et al. 2026). Although scalable, these explanations are constrained to existing annotation categories, are derived from correlations with a large corpus of proteins rather than a protein-specific role, and are applied to SAE latents selected based on activation patterns, not their causal role.

### Sparse feature circuits and activation patching

Mechanistic interpretability in NLP has converged on two complementary tools for causal analysis: *activation patching* (intervening on specific components and measuring the effect on the output) (Olah et al. 2020) and *attribution methods* such as integrated gradients (Sundararajan, Taly, and Yan 2017). Marks et al. unified these ideas in the *sparse feature circuits* framework, treating SAE features as nodes in a computational graph and ablating values in that graph to identify the features causally responsible for a task-specific prediction from a single sequence (Marks et al. 2025). Steering similarly modifies SAE activations, but serves a different purpose: whereas activation patching is used to identify causal features, steering uses selected internal features to alter model behavior. It can therefore provide complementary validation of causally identified features, although steering alone can be applied to arbitrary features and does not establish their causal role in the original computation. Hence, we adapt the sparse feature circuits framework to zero-shot mutation effect prediction in ESM-2, introducing a Jensen-Shannon recovery score suited to the masked-marginal scoring setting and performing the first mechanistic circuit analysis of an unsupervised mutation-effect prediction in a pLM.

## 3 Methods

### Zero-Shot Mutation-Effect Prediction Formulation

Following Meier et al. (2021), we score mutation effects using the masked-marginal log-likelihood ratio (LLR). Given a wild-type sequence *x*^wt^ and a mutation at position *t* from 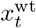 to 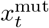, let 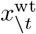 denote the sequence with position *t* masked. The mutation-effect score of ESM-2 650M is defined as

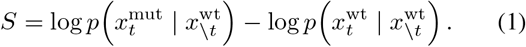

This score measures whether the model assigns higher probability to the mutant or wild-type residue using only the surrounding sequence context.

### Models and Data

We use the SAEs trained by Adams et al. (2025) on the residual stream of ESM-2 650M at multiple layers to identify features contributing to the model’s zero-shot mutation-effect predictions. We refer to these SAE features as *latents* throughout this study. Additional details on the base model and SAE checkpoints are provided in Appendix A.

We select a case-study protein from the ProteinGym single-substitution DMS benchmark (Notin et al. 2023). We rank proteins by the zero-shot Spearman correlation between ESM-2 650M masked-marginal scores and DMS measurements, and select `DNAJA1_HUMAN` (*ρ* = 0.803), one of the highest-scoring proteins, because it contains a well-characterized and short 67-residue J-domain protein, making it tractable for residue-level feature analysis. We choose L7D as the primary mutation because it is among the most deleterious substitutions in the assay and is also strongly scored as deleterious by ESM-2 650M. Biochemically, replacing Leu7 with aspartic acid introduces a charged side chain into a hydrophobic local environment, providing a plausible hypothesis about the destabilizing mechanism against which we can evaluate recovered features.

### Selection of Mutation Relevant Latents

#### Ranking latents by causal effect

We identify mutation-relevant SAE latents using the *single-input zero-ablation* procedure from sparse feature circuits (Marks et al. 2025).

For an input sequence **x**_*\*t_ where the mutated residue at position *t* is masked, let 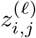 denote latent *j* at token position *i* in layer *l*. We quantify the causal importance of each latent– token node using integrated gradients (Sundararajan, Taly, and Yan 2017) to estimate it’s zero-ablation indirect effect (Pearl 2022) on the mutation-effect score *S*:

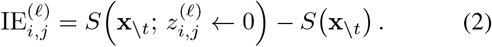

Here, 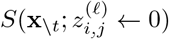 denotes the mutation-effect score obtained after setting the selected SAE latent–token node to zero and continuing the intervened computation through ESM-2 650M. Let 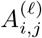 denote the resulting integrated-gradients attribution. We then rank latent–token nodes by 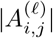, prioritizing those estimated to contribute most strongly to the zero-ablation indirect-effect target. We use zero rather than mean ablation because zero represents the inactive state of a Top-*K* SAE latent. As a baseline, we also rank latents by activation difference, following Wang et al. (2026). Further details provided in Appendix C.

#### Recovery score formulation and circuit selection

After ranking latent–token pairs by their indirect effect, we evaluate how much of the original model computation can be recovered by a selected subset of causally relevant pairs, which we call a circuit *C*. For a fixed layer *l*, let *M* denote the complete circuit where all SAE latent-token pairs are kept at their original activations. Let ∅ denote the empty circuit, where all candidate nodes are replaced by their dataset mean activations at the latent-token level. These means are estimated from forward passes over 10,000 randomly sampled UniRef50 proteins. For a recovered circuit *C*, nodes in *C* are restored to their observed activations, while all remaining candidate nodes are mean-ablated.

We evaluate recovery at the masked mutation position *t* using two metrics: The first, the Log-Likelihood Ratio (LLR) recovery score, whichmeasures recovery of the mutation-specific log-likelihood ratio relative to the empty-circuit baseline (Appendix D.3) at the masked position *t*. The second measures recovery of the model’s complete output distribution over the 20 standard amino acids at the masked position. Let *P*_*M*_, *P*_*C*_, and *P*_*∅*_ denote the output distributions under the clean computation, recovered circuit, and empty circuit, respectively. We define the Jensen–Shannon recovery score as

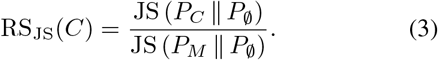

Intuitively, this score measures the fraction of the clean model’s distributional shift away from the mean-ablated baseline that is recovered by circuit *C* and is used as the primary method of recovery in all the downstream analyses. A score of 1 indicates that the selected circuit recovers the clean output distribution, whereas a score of 0 indicates no improvement over the empty circuit. We construct the recovered circuit by greedily adding latent–token pairs in decreasing order of 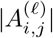, and select the smallest circuit that reaches the recovery threshold *θ* = 0.8. The empty circuit construction and full circuit recovery procedures are described in Appendix D.1 and D.2. The denominator of our recovery score can, in principle, approach zero if the empty circuit is nearly identical to the clean circuit. In our experiments, this does not occur: across different empty-circuit baselines, the Jensen–Shannon divergence from the clean distribution remains well separated from zero.

#### Latent interpretation with existing tools

To interpret the constituent latents of the selected latent–token circuits, we adapt ToolUniverse’s variant-interpretation tools to our corresponding ESM-2 650M SAE latents. Tool details are provided in Appendix G. We additionally inspect InterProt visualizations (Adams et al. 2025) to examine each latent’s positional activation pattern and guide manual annotation.

#### Structure-guided, mutation-specific latent interpretation

To complement the corpus-level annotations obtained from ToolUniverse and InterProt, we perform a structure-guided analysis of the selected DNAJA1 circuit latents. For each SAE layer, we select the top 10–15 latents ranked by 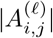^)^ and visualize in PyMOL version 3.1.8 (Schrödinger, LLC 2026) the residue positions at which each latent contributes to the circuit, relative to the wild-type residue at the mutation site. We then assess whether these positions exhibit plausible sequence or structural relationships with the mutation site and, where available, compare the observed patterns with prior biochemical or structural evidence. Otherwise, we evaluate whether the pattern is consistent with a plausible mechanism for the predicted mutation effect.

#### Circuit latent reuse across same-position mutations

We next extend the mutation-specific analysis to ask whether substitutions at the same residue recruit common SAE representations and whether the reused latents encode shared biochemical or structural patterns. We consider four mutation families, `L7X`, `L51X`, `H30X`, and `L27X`, where `X` denotes the substitutions assayed at that position in the corresponding ProteinGym DNAJA1 DMS dataset (Notin et al. 2023). Note that while there are 76 possible substitutions at these four sites, we only analyze the 67 included in ProteinGym. These families represent two distinct structural contexts and effect severities: `L7` and `L51` are buried positions with generally strongly deleterious substitutions, whereas mutations at `L27` and `H30` tend to have more moderate effects; `H30` additionally lies within the conserved HPD motif of the J-domain in DNAJA1. For each mutation, we select the circuit reaching recovery threshold *τ* = 0.8 by ranking latent–position pairs according to 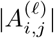. To quantify circuit reuse, we compare the sets of unique SAE latent IDs represented in each pair of mutation-specific circuits for each mutation family using Jaccard similarity. Latents appearing in both circuits are treated as shared latents. We interpret these shared latents using the same structure-guided, mutation-specific procedure described above.

## 4 Recovery Trends Across Mutations

### IE-based ranking yields higher and sparser recovery efficiency

We compare the four combinations of ranking method and recovery score for their ability to recover a sparse computational circuit for the 67 mutations at `L7`, `L51`, `H30`, and `L27` (Figure 2). For each mutation and each layer of SAEs, we seek to compute a resolvable circuit, defined as a set of latent-token pairs at a layer that, when patched into a mean-ablated circuit, achieve the 0.8 recovery threshold. We then compare the number of latent-token nodes in the resolvable circuit. For computational feasibility and to ensure the recovered circuits are interpretable, we patch only the first 10, 000 latent-token pairs for each circuit.

**Figure 1:**
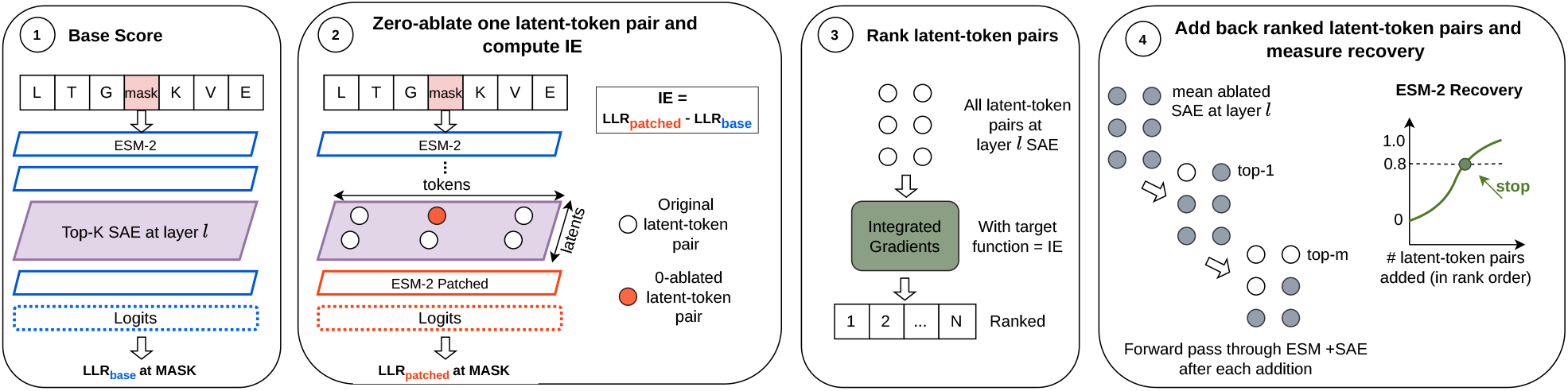
Overview of the sparse feature-circuit discovery framework. For each layer with trained SAEs, after computation of a baseline score, zero-ablation activation patching is used to compute the Indirect Effect, a measure of causal influence, of each latent-token pair. We then restore latent-token pairs in order of causal influence to a fully ablated model, stopping when we reach a designated threshold of recovery.

**Figure 2:**
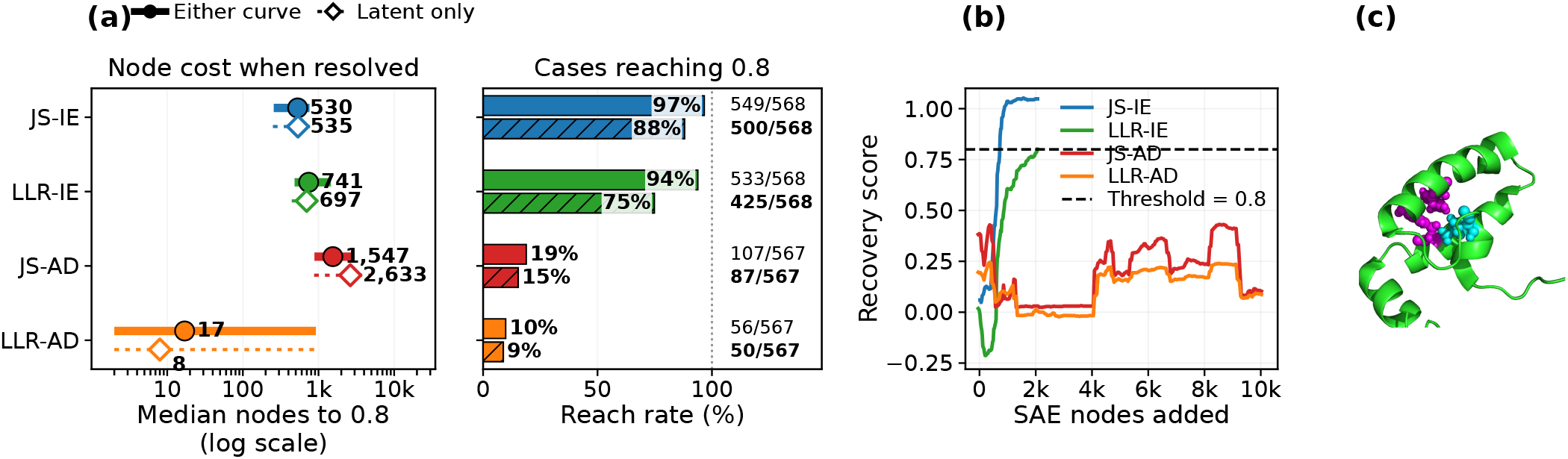
Recovery efficiency and structural interpretation of SAE circuit latents. (a) Median circuit size among resolved mutation–layer cases (left) and fraction of cases reaching the *τ* = 0.8 recovery threshold (right). Resolved represent the cases reaching the threshold. Solid markers and bars report the earlier threshold crossing across the latent-only and latent-plus-reconstruction error node recovery curves; open markers and hatched bars report recovery with latent-only nodes. AD-ranked cases are counted as resolved only when the threshold is reached within 10,000 nodes. (b) Layer-16 recovery trajectories for the representative `L7D` mutation across different recovery strategies. (c) DNAJA1 J-domain structure (PDB `2LO1`, chain A) (Dutta et al. 2012) showing the mutation site L7 in cyan and circuit-associated residues Y22 and Y48 in magenta, identified among the five highest-ranked JS–IE latents.

We observe that Indirect Effect (IE) ranking recovers the model output far more reliably than Activation Difference (AD), and that the Jensen-Shannon (JS) recovery score produces smaller circuits than the log-likelihood ratio (LLR) score. JS-IE and LLR-IE resolve 88% and 75% of circuits, with medians of 535 and 697 nodes in the circuit, respectively. JS-AD and LLR-AD recovery resolve only 15% and 9%, with medians of 2,633 and 17 nodes in the circuit, respectively. We note that the small median circuit size for LLR-AD is computed from less than 10% of mutations, and the remaining majority were not resolvable even with 10,000 nodes. We repeat this analysis while including the SAE reconstruction error nodes, and find that the same trends hold but a moderately higher fraction of circuits are recoverable when error nodes are included (*e*.*g*., 97% of circuits now resolvable using JS-IE, versus 88% previously). For more details, see Appendix F.1.

The circuits computed with JS-IE and LLR-IE use only approximately 0.20% and 0.28% of the 266,112 candidate latent-token nodes in a layer on an average, showing that the mutation-effect prediction can be recovered through a highly sparse subset of SAE activity at a layer. The corresponding interquartile ranges are 256-759 nodes for JS-IE and 483-1,457 nodes for LLR-IE, indicating that JS recovery produces consistently smaller circuits across mutations. Further details on reliability of JS-IE based recovery are provided in Appendix F.2 and F.3.

## 5 Interpretation of Circuits

### JS-IE circuits recover nonlocal structural and functional context

Having established the recovery efficiency of JS-IE, we next examine the biological context represented by its highest-ranked circuit nodes. We focus on layer 16 because the aggregate analyses identify it as a representative depth with substantial circuit reuse and clear enrichment of contact-associated latents (Figure 4(a, b)). For the representative `L7D` mutation, Figure 2(c) shows that the circuit prioritizes `Y22` and `Y48`, which are distant from `L7` in sequence but lie within its three-dimensional contact neighborhood, defined by a 5 Å side-chain heavy-atom cutoff. Given that the DMS assay measures folding stability, this pattern is consistent with a hypothesis in which replacing the buried hydrophobic leucine with a charged aspartate perturbs local packing interactions involving these residues. More broadly, of the 87 circuit latents (Figure 3), contacts comprise the largest category (39.1%), followed by the HPD motif (10.3%), a signature of the J-domain proteins which promotes Hsp70 ATPase activity (Tsai and Douglas 1996; Fan, Lee, and Cyr 2003; Kampinga and Craig 2010), and other residues in the surrounding Hsp70-interaction surface (9.2%) (Greene, Maskos, and Landry 1998; Kityk, Kopp, and Mayer 2018). Amino-acid or sequence-pattern detectors contribute another 9.2%, with the remainder spanning local, structural, functional, and unresolved categories. Thus, the `L7D` circuit combines mutation-site packing information with broader protein-specific sequence and functional context. Results for ranking–recovery combinations for other mutations are provided in Appendix H.1.

**Figure 3:**
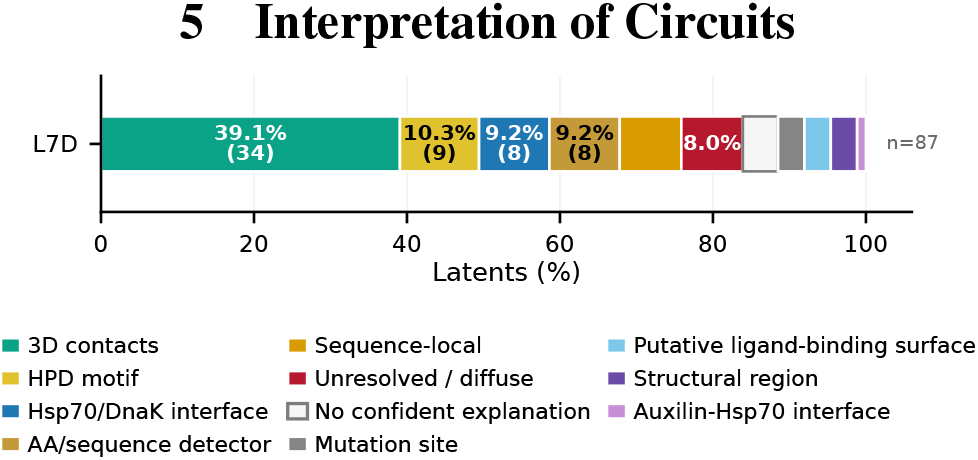
Layer-16 latent composition for the representative `L7D` mutation. Colors indicate the biological interpretation assigned to each latent.

**Figure 4:**
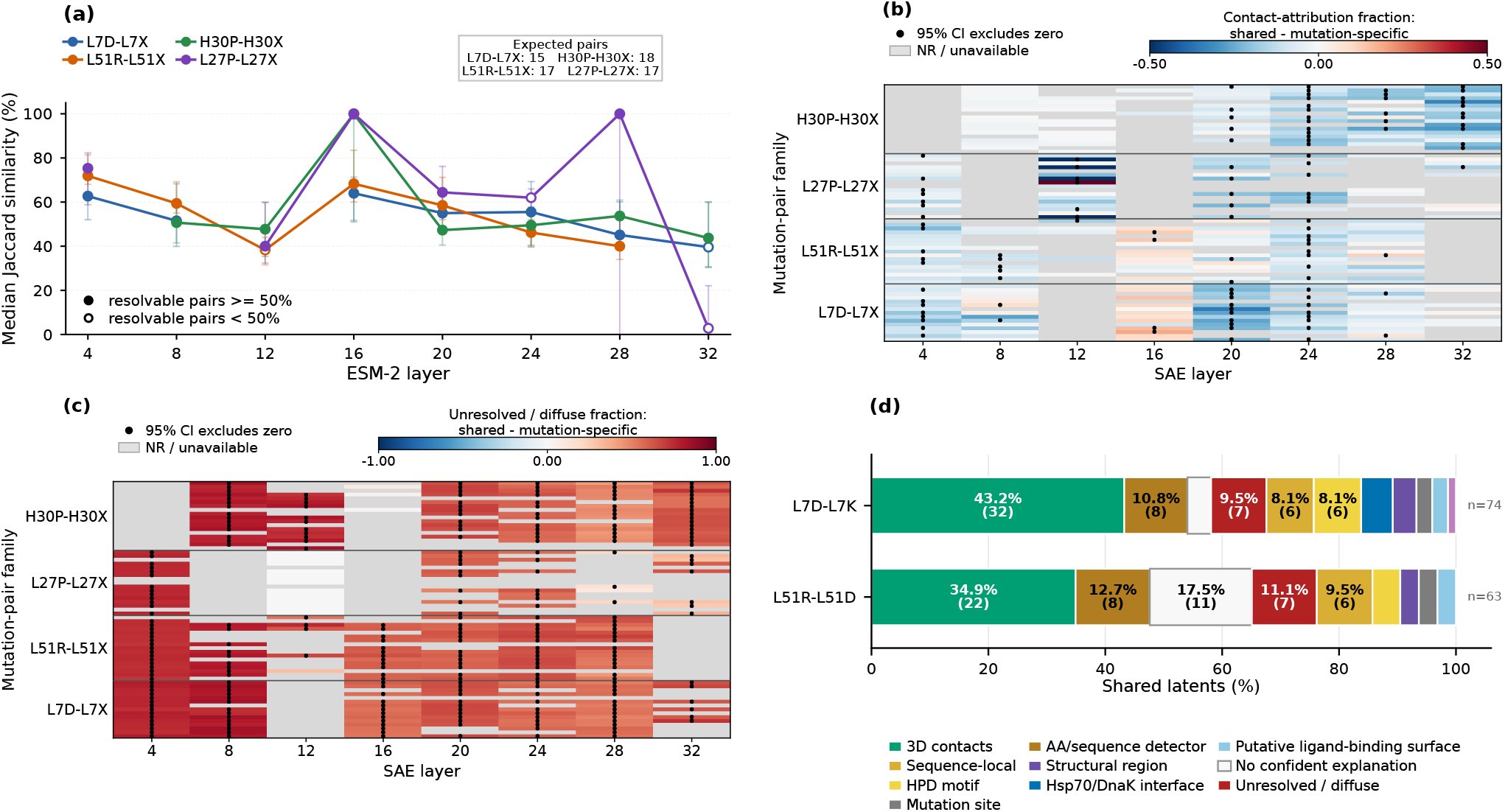
Analysis of shared mutation-relevant circuits. **(a)** Family-level median circuit similarity across model layers at the *τ* = 0.8 recovery threshold. Filled markers indicate layers for which at least 50% of the expected mutation pairs are resolvable; open markers indicate lower resolvable fraction. A mutation pair is resolvable when both mutations reach the *τ* = 0.8 recovery threshold within the 10,000-node cap. **(b)** Difference in three-dimensional contact-attribution fraction between shared and mutation-specific circuits across SAE layers and mutation families. **(c)** Corresponding difference in unresolved/diffuse attribution fraction. In **(b)** and **(c)**, positive values indicate enrichment in shared circuits, negative values indicate depletion, black points mark comparisons whose 95% confidence intervals exclude zero, and gray cells denote unavailable comparisons. The color range in (b) is set within ±0.5 (as opposed to (c)) to clearly illustrate the contact enriched cases. **(d)** Manual annotation composition of layer-16 shared latents for the `L7D-L7K` and `L51R-L51D` mutation pairs.

### Mechanistic Descriptions of Mutation-Effect Circuits using ToolUniverse SAE Agents

Having identified mutation-specific structural and functional patterns in the `L7D` circuit, we next ask whether corpus-based SAE interpretation methods highlight the same mechanisms. We adapt ToolUniverse’s variant-effect feature-extraction and labeling pipeline to our layer-specific ESM-2 SAE check-points. Seven of the ten highest-ranked `L7D` latents received the broad label *secondary-structure associated*. Although not incorrect, this description does not explain the residue-contact context through which these latents contribute to the predicted deleterious effect as discussed above. Only two latents were labeled *structural-stability associated*, while another was labeled *protein-domain associated* despite showing no overlap with annotated domain features in DNAJA1. InterProt annotations (Adams et al. 2025), which are likewise derived from corpus-level activation patterns, also diverged from the latents’ apparent mutation-specific roles in all the cases. Examples of these divergent concept interpretations are provided in Appendix H.2.

### Biological Annotation Categories

Because ToolUniverse and InterProt did not reliably capture mutation-specific context, we define six scalable categories for consistently annotating latents across mutations based on their residue positions and structural relationships to the mutation site in DNAJA1 structure. These latent categories are: **mutation site**, activating only at the mutation position; **contacts**, activating on tokens representing residues whose side-chain heavy atoms lie within 5 Å of the mutation site; **sequence-local**, activating on tokens within a ±5-residue window around the mutation site; **HPD motif**, activating at positions 30–32; **other residues**, localized patterns not covered by the preceding categories; and **unresolved/diffuse**, broadly distributed patterns without a coherent structural, sequence-local, motif, or residue-identity interpretation. To avoid assigning specific interpretations to diffuse patterns, we restrict definite annotations to latents localized to at most five token positions, a practical threshold for the short DNAJA1 J-domain fragment. Detailed assignment criteria are provided in Appendix H.3.

### Shared Representations Across Mutations

#### Within-family mutations reuse substantial latents but in a layer-dependent manner

With the scalable annotation framework in place, we first quantify how much of the recovered latent set is reused across substitutions at the same residue before examining the biological context encoded by the shared latents. As shown in Figure 4(a), median Jaccard similarities fall between approximately 40% and 75% for most resolvable family–layer combinations, indicating substantial rather than disjoint circuit reuse. The `L7D-L7X` and `L51R-L51X` families exhibit comparatively stable, moderate sharing across depth, whereas `H30P-H30X` and `L27P-L27X` are more layer selective. Both reach 100% median similarity at layer 16, and `L27P-L27X` again reaches 100% at layer 28 before dropping sharply at layer 32; this should be interpreted cautiously because fewer than half of the expected pairs are resolvable. Thus, circuit reuse is substantial overall but depends on both mutation family and model depth. Circuit-sharing distributions are provided in Appendix I.1.

#### Shared latents are contact enriched in preferred layers and mutation-families

Having observed substantial circuit sharing within mutation families, we next ask whether the shared nodes encode common biological information relevant to mutation-effect prediction. For each annotation, we define Δ*F* as the difference between the fraction of |*A*| attribution in the shared circuit and that in the corresponding mutation-specific components; positive values indicate enrichment in the shared circuit. Markers denote comparisons whose 95% confidence intervals exclude zero, while grey cells indicate that one or both circuits did not reach the recovery threshold. Among the biological annotations considered, contacts show the clearest positive enrichment at selected layers (Figure 4(b)), whereas most other categories are predominantly depleted (Appendix I.3). We therefore first examine how this contact-associated signal varies across mutation families and layers. Contact enrichment is most evident for the buried `L7X` and `L51X` families, but is depth-dependent. At layer 16, positive enrichment occurs in approximately one quarter of resolvable pairs in both families (23% for `L7X` and 25% for `L51X`), whereas other layers show weaker or reversed trends. The more weakly deleterious `L27X` and `H30X` families exhibit less consistent contact enrichment.

Interestingly, the largest portion of shared-circuit attribution is significantly enriched (as shown by markers) in the unresolved/diffuse category across many resolved pair-layer comparisons (Figure 4(c)). These latents distribute attribution across multiple positions without aligning with any predefined annotation. We therefore examined the layer-16 diffused latents from `L7D-L7K` and `L51R-L51D` using InterProt (Adams et al. 2025) and found that they generally activate broadly across protein sequences. In two cases, corresponding to latent IDs `3556` and `546`, the InterProt examples included activation across the J-domain of DNAJA1 or another J-domain-containing protein, suggesting possible sensitivity to domain-level context. Detailed enrichment calculations are provided in Appendix I.2.

#### Shared-circuit compositions are contact-centered but encode diverse concepts

To complement the enrichment analysis, we manually annotate the layer-16 shared latents from two illustrative pairs at buried mutation sites, `L7D-L7K` and `L51R-L51D` (Figure 4(d)). We select these pairs because their families show strong contact enrichment at layer 16 and their shared circuits contain sufficiently many interpretable latents for detailed analysis. Contacts form the largest category, accounting for 43.2% and 34.9% of shared latents, respectively. Both circuits also contain amino-acid or sequence-pattern detectors (10.8% and 12.7%), together with sequence-local, HPD-motif, and other protein-specific interaction patterns. Thus, the interpretable shared computation combines mutation-specific context, including local structural contacts and context-dependent residue identity, with protein-level features such as conserved motifs and putative ligand-binding surfaces. This suggests that substitutions at the same position are evaluated by the model using both their local structural consequences and broader properties of the candidate protein scaffold.

## 6 Discussion and Conclusion

We introduce a sparse feature-circuit dicovery framework for identifying the SAE latents causally involved in zero-shot mutation-effect prediction. Our causal indirect-effect (IE) ranking resolves compact circuits more reliably than non-causal activation difference based selection, which is currently implemented in many leading tools. We observed that activation differences do identify causally relevant nodes early for a small subset of mutations (≈ 9%), but they do not consistently prioritize the features that mediate unsupervised mutation-effect prediction. This indicates that causal interventions are necessary to reconstruct network behavior. Among IE-ranked circuits, we find that our Jensen-Shannon recovery score produces smaller circuits than the mutant-residue probability changes measured by LLR.

When attempting to annotate our causally relevant latents with biological function, we found that existing tools provided only general explanations that did not match our detailed observations about the protein structure, failing to recognize that the highest important latents corresponded to contacts of the mutated site. However, existing tools were able to explain certain global patterns — such as latents that activate on the presence of the DNAJA1 domain — that were missed by our protein-specific explanations. This indicates that future development of latent interpretation agents should combine analysis of protein-specific roles with global roles from across a large corpus of sequences.

We observe notable patterns of latent reuse across circuits. In particular, for substitutions at buried residues, latents that appear to encode residue-residue contact information are preferentially reused at layer 16. This greater reuse may arise because buried sites impose conserved packing constraints in the three dimensional environment, enabling the model to reuse contact-associated features across substitutions for predicting mutation effects. Latents that encode diffuse signal are also enriched among shared latents. We found that in some cases this diffuse signal does represent a protein domain detector for relevant proteins, though in other cases it was not interpretable. The enrichment of diffuse latents in shared circuits may therefore reflect reusable, protein-level contextual features that complement the more localized mutation-specific context necessary for the mutation-effect prediction at a given site.

We find that even on a small, well-characterized protein where ESM-2 650M is successful at unsupervised mutation effect prediction, a substantial amount of network behavior remains uninterpretable. Only 88% of our studied mutations and layers have a resolvable latent-only circuit, and we could provide a plausible biological explanation for at most 90% of latents, in even the most interpretable circuit. This will likely improve with better protein language models, better mechanistic interpretability methods, and better latent interpretation methods.

## Limitations

Our study is limited to ESM-2 650M and a single protein, with detailed interpretation concentrated at one SAE layer and two representative mutation pairs. Accordingly, the observed circuit reuse and contact-centered organization may depend on the DNAJA1 fold or the selected pLM and SAE representation. Future work will extend unsupervised mutation effect prediction to more proteins and models. Moreover, substitutions at the same position share an identical masked input, so their overlap reflects a common contextual computation in addition to any shared biochemical mechanism.

### Conclusions

Our work provides the first causal circuit analysis for unsupervised mutation effect prediction in a pLM, and points to clear next steps for the application of mechanistic interpretability to pLMs. First, we suggest the use of IE rather than activation difference to prioritize latents where feasible, including for steering network behavior. Second, we suggest that existing latent annotation tools be improved to include protein-specific explanations. The tools for circuit reconstruction introduced in our work are readily applicable to other proteins: latent ranking methods, and recovery scores. Our metrics — percent of circuits resolvable, circuit size, and percent of latent-token pairs that are interpretable — could facilitate head-to-head comparison of mechanistic interpretability methods of pLMs.

## Appendix

### A Models

#### Protein Language Model

We use ESM-2 650M (Lin et al. 2023) as our base model for all experiments.

#### Sparse autoencoders (SAEs)

Neural-network representations can be polysemantic, with individual neurons encoding multiple unrelated features, potentially due to superposition (Olah et al. 2020; Bricken et al. 2023). SAEs aim to disentangle these dense representations into sparse, more interpretable latent features (Elhage et al. 2022). We use the publicly available Top-*K* SAEs of Adams et al. (2025), trained on residual-stream activations. For an activation *x* ∈ ℝ^d^, the SAE computes

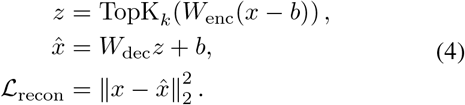

Here, *W*_enc_ and *W*_dec_ are the encoder and decoder matrices, *b* is a bias term, and only the *k* largest latent activations are retained. The reconstruction objective preserves information from the original representation, while Top-*K* sparsity encourages individual latent dimensions to respond to a limited set of patterns. These approximately monosemantic latents provide interpretable units for mechanistic analysis and causal intervention (Gao et al. 2025a).

#### Parameters used for the SAE model

We use the pretrained top-*k* SAEs of Adams et al. (2025) (InterProt), one per residual block of ESM-2-650M (`esm2_t33_650M_UR50D`). Our experiments use the layer-16 SAE (`esm2_plm1280_l16_sae4096`) on the frozen backbone.

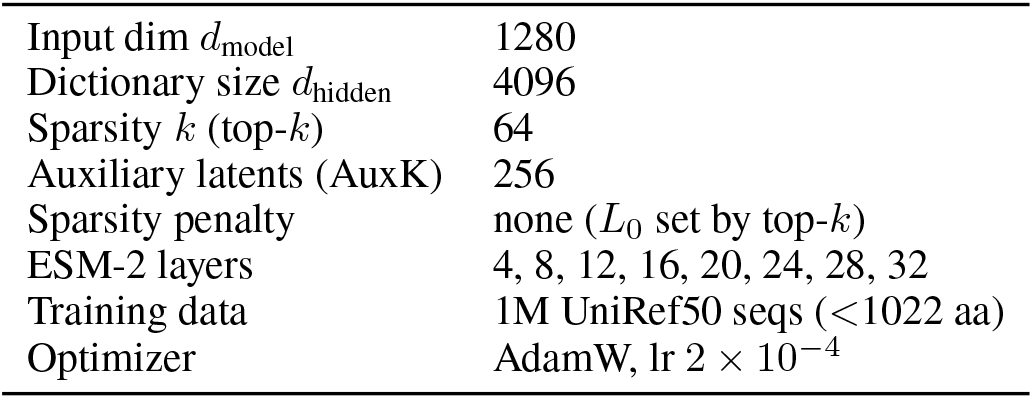

|  |  |
| --- | --- |
| Input dim $d_{\text{model}}$ | 1280 |
| Dictionary size $d_{\text{hidden}}$ | 4096 |
| Sparsity $k$ (top-$k$) | 64 |
| Auxiliary latents (AuxK) | 256 |
| Sparsity penalty | none ($L_0$ set by top-$k$) |
| ESM-2 layers | 4, 8, 12, 16, 20, 24, 28, 32 |
| Training data | 1M UniRef50 seqs (< 1022 aa) |
| Optimizer | AdamW, lr $2 \times 10^{-4}$ |

### B Experimental Setup Hardware

All experiments were run on the UMass Unity cluster, using a single NVIDIA GTX 1080 Ti GPU per job (8 CPU cores, 80 GB RAM). The full pipeline–baseline scoring, latent ranking, and circuit-recovery analysis on DNAJA1 and 67 mutations–took approximately 41 mins on the GPU per mutation per layer run on average when circuit discovery is run across 8 layers for 67 mutations each with circuit discovery times reaching upto 3 hours for weakly deleterious mutations like H30X.

### C Ranking Latents by Activation Difference

As a non-causal baseline, we rank latent–token nodes by the difference between their activations on the unmasked mutant and wild-type sequences, following Wang et al. (2026). Let **x**^mut^ and **x**^wt^ denote these sequences. For latent *j* at token position *i* in layer *l*, we compute

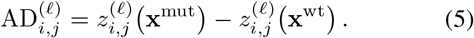

We rank nodes by 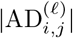, prioritizing features whose activations change most between the mutant and wild-type inputs. Unlike IE ranking, this method measures representational change without intervening on the model computation.

### D Circuit Construction and Recovery

#### D.1 Empty Circuit Construction

The empty circuit ∅ is the baseline with which all recovery changes are measured. It is first found by replacing every candidate latent–token node with its dataset-mean activation, which was estimated from forward passes over 10,000 randomly sampled UniRef50 proteins using reservoir sampling.

##### Algorithm 1

Constructing the empty circuit

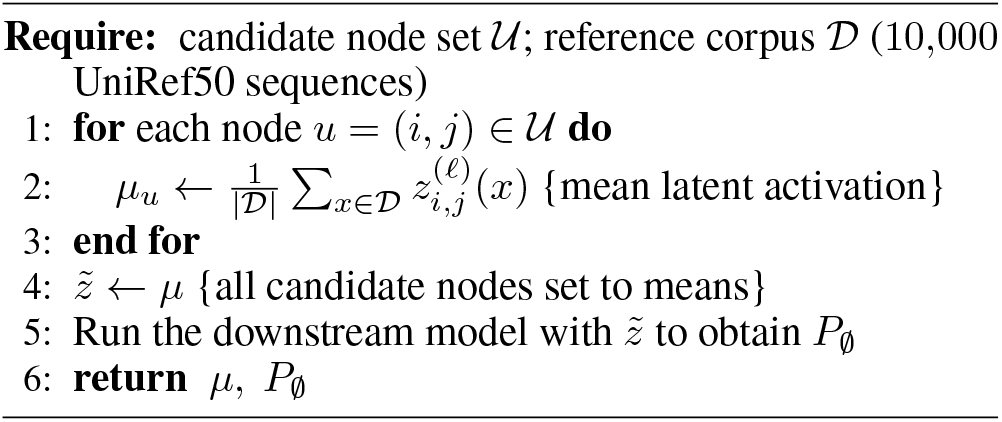

##### Algorithm 2

Constructing the recovered circuit

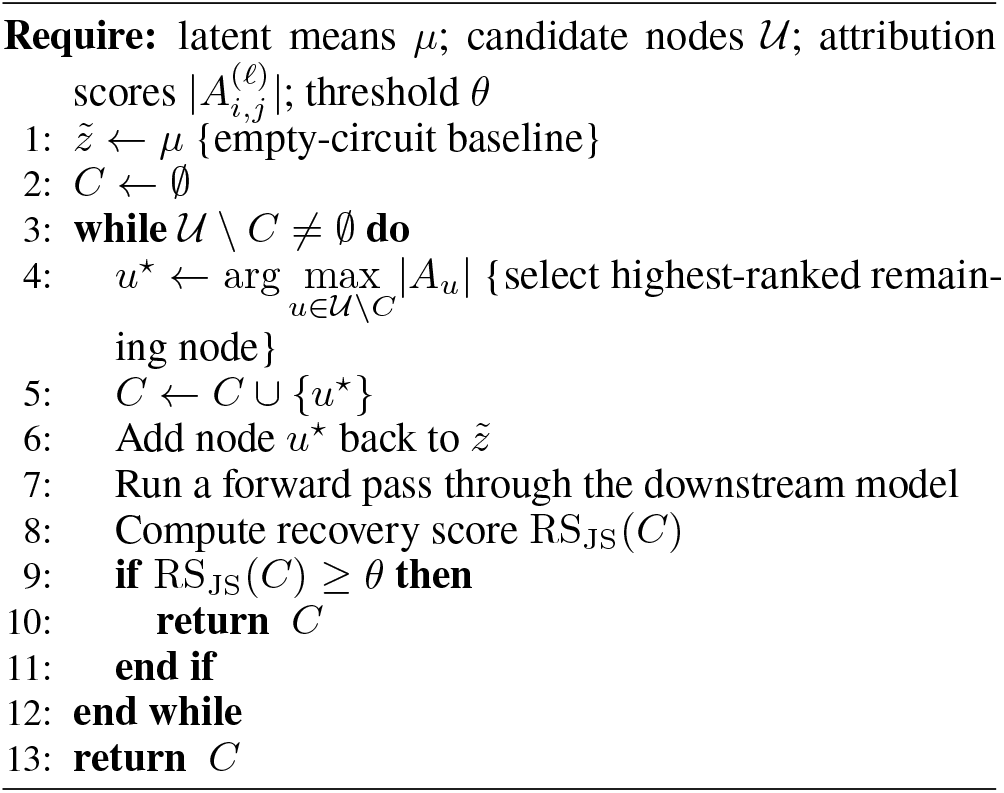

This baseline preserves generic protein information, yielding an approximately uniform output distribution at the masked position that is well separated from the clean distribution. The algorithm for empty circuit construction is described in Algorithm 1.

#### D.2 Circuit Recovery Algorithm

We construct a recovered circuit by running a forward pass through the top ranked latent–token nodes added back greedily in decreasing order of 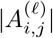 while keeping all remaining candidate nodes at their empty-circuit or mean values, then measuring the output distribution. The computation stops with the smallest circuit reaching the recovery threshold *θ* = 0.8, as described in Algorithm 2.

#### D.3 Log-likelihood Ratio (LLR) Recovery Score

Following the setup in Recovery score formulation and circuit selection section, we define the recovery score of a candidate circuit *C* in general as:

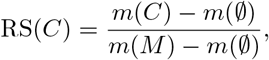

where *m*(·) represents a chosen measure of circuit behavior as per (Marks et al. 2025).

Adapting this method to our case, we use the masked-marginal score *S* (Eq. 1) directly at the masked residue position *t* we define LLR recovery score as:

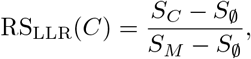

where *S*_*M*_, *S*_*C*_, and *S* are the masked-marginal scores under the clean, recovered, and empty circuits. This formulation is used to measure the recovery of just the scalar masked-marginal mutation-effect score, while RS_JS_ measures recovery of the entire distribution over all 20 amino acids.

### E Empty-Circuit Sensitivity

The empty circuit defines the reference distribution against which all recovery scores are measured. We therefore assess whether its construction is sensitive to the dataset used to estimate the mean SAE latent activations and reconstruction errors. For each construction, we run the mean-ablated model and examine its probability distribution over the 20 standard amino acids at the masked mutation position *t*. Similar distributions across datasets would indicate that the empty-circuit baseline is robust to the choice of reference data.

We compare two empty-circuit constructions, shown in Figure 5:

**Figure 5:**
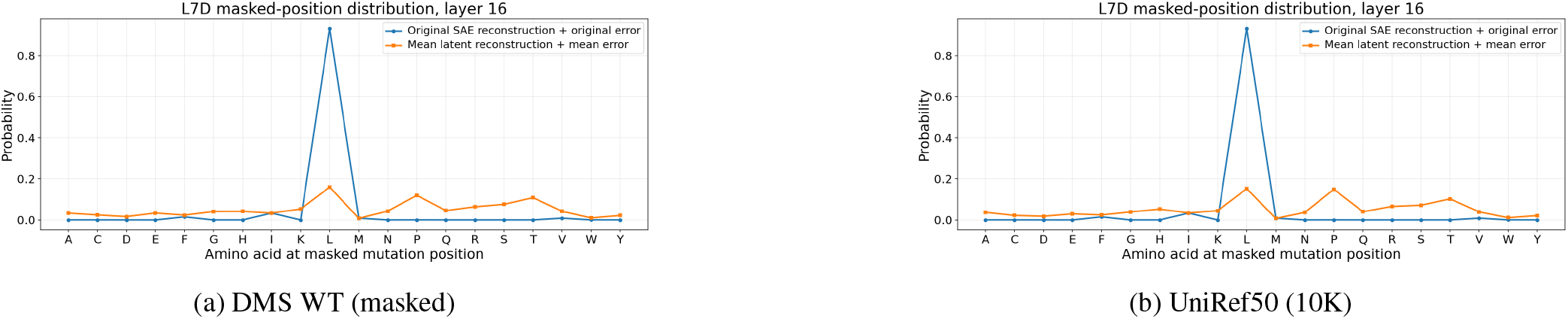
Empty circuit baseline comparison for L7D at layer 16. Each panel shows the probability distribution at the masked mutation position under the original SAE reconstruction (blue) versus the empty circuit constructed from different mean activation datasets (orange). (a-b) Broader baselines produce approximately uniform empty-circuit distributions, leading to well separated recovery denominators. The baselines are visually similar, indicating the stability of mean activation structure on diverse protein datasets.

**Figure 6:**
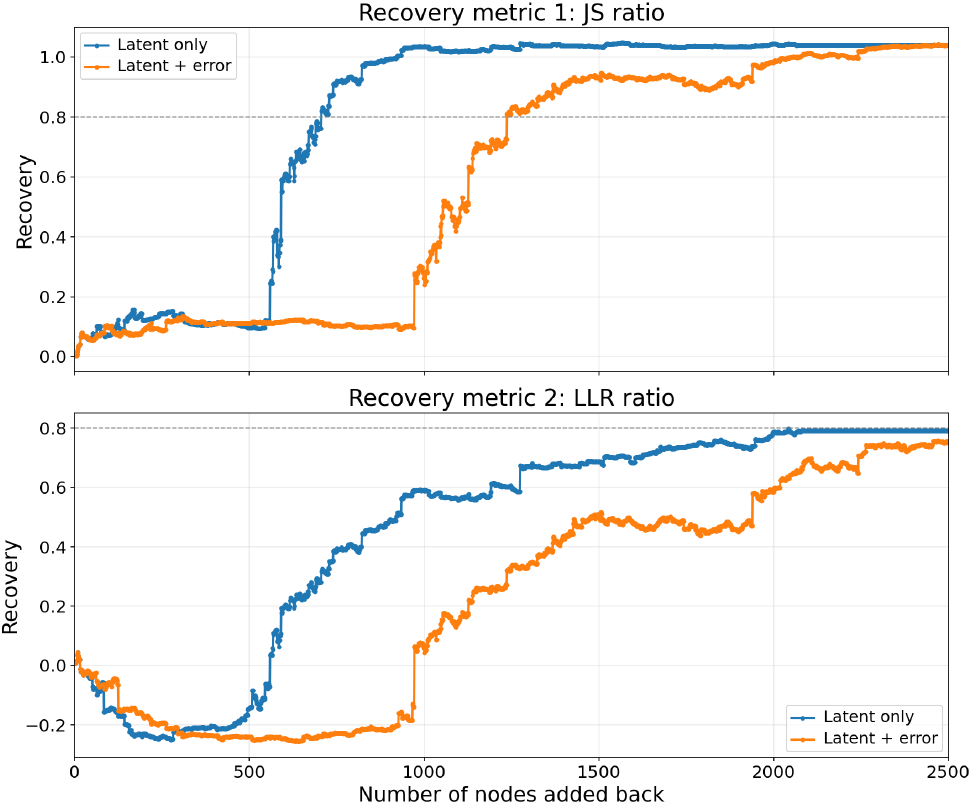
Latent-only versus latent-and-error recovery for L7D at layer 16. JS-ratio (top) and LLR-ratio recovery (bottom) is shown as a function of the IG-ranked nodes restored, with a *τ* = 0.8 threshold indicated by the dashed line. Both panels show including SAE reconstruction-error nodes (orange) allows for the same recovery level as the latent-only circuit (blue), but takes much longer to cross the threshold since including the error term spreads the recovery signal across more components.

- **Masked ProteinGym wild-type sequences:** Mean latent activations and reconstruction errors are computed over the wild-type sequences from ProteinGym single-substitution DMS assays, with each sequence position masked in turn to match the input context used for masked-marginal scoring (Figure 5a).
- **Random UniRef50 sample:** Mean latent activations and reconstruction errors are computed over 10,000 UniRef50 sequences selected using reservoir sampling, with each sequence position again masked providing a broader sample of the protein activation landscape. The resulting empty-circuit distribution (Figure 5b) is nearly identical to the ProteinGym-based distribution, indicating that the estimated mean activation structure remains stable when computed from sufficiently large and diverse protein datasets.

### F Recovery Trends Across Mutations

#### F.1 Latent-Only vs. Latent-and-Error Recovery

An SAE decomposes each model activation *x* into its reconstruction 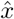 and the residual 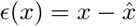. Following Marks et al. (2025), we represent this residual at each token position as a *reconstruction-error node*. Unlike SAE latents, error nodes are not individually interpretable; they retain information in the original model activation that is not captured by the SAE dictionary. We repeat circuit recovery with these error nodes added to the candidate latent–token nodes and rank all candidates using the same attribution criterion. This allows us to distinguish behavior recovered by interpretable SAE latents from behavior that additionally depends on information remaining in the reconstruction error.

Including error nodes increases the fraction of mutation– layer cases reaching the 0.8 recovery threshold. For IE ranking in Figure 2, the reach rate increases from 88% to 97% under JS recovery and from 75% to 94% under LLR recovery; the corresponding AD increases are smaller, from 15% to 19% and from 9% to 10%, respectively. Thus, most IE-ranked cases can be recovered using SAE latents alone, while the reconstruction error retains substantial mutation-relevant information in cases where latent-only recovery plateaus below the threshold.

#### F.2 JS-Divergence Trajectories

Figure 7 reports the Jensen–Shannon divergence between the output distributions of the original model *M* and the recovered circuit *C*. This provides a direct validation of the recovery metric: as circuit recovery approaches 1, the divergence should approach 0, indicating that the recovered circuit reproduces the original amino-acid probability distribution. This confirms that the added latent–token nodes restore the mutation-relevant signal rather than merely increasing the recovery score. For the buried, strongly deleterious mutation L7D (Figure 7a), the latent-only JS divergence begins near the empty-circuit baseline (≈ 0.5) and holds there while the earliest-ranked nodes are restored, then drops in the 550–850 node range and converges to near zero within around 1,000 nodes. The horizontal axes are not shared across panels as L7D spans ~ 2,500 restored nodes, while L51R, H30P, and L27P (Figure 7b– d) extend to 10, 000. L51R (b) is plotted on this wide axis but, like L7D, reaches near-zero divergence after only a small number of restored nodes, which appears as a flat line at zero, and shows that few latents are required for full recovery. The more weakly deleterious H30P and L27P (c–d) share the initial decline but retain some residual divergence for the latent-only case across the plotted range rather than converging to zero, reflecting the lower resolvability of weakly deleterious families. However, for both mutation families, the divergence approaches zero when reconstruction-error nodes are included, even when latent-only circuits remain incomplete. This indicates that, in cases not fully captured by the SAE latents, the reconstruction error retains a substantial portion of the mutation-relevant information, specially for hard mutations where it is hard to estimate the exact reasons for observed mutation effects.

**Figure 7:**
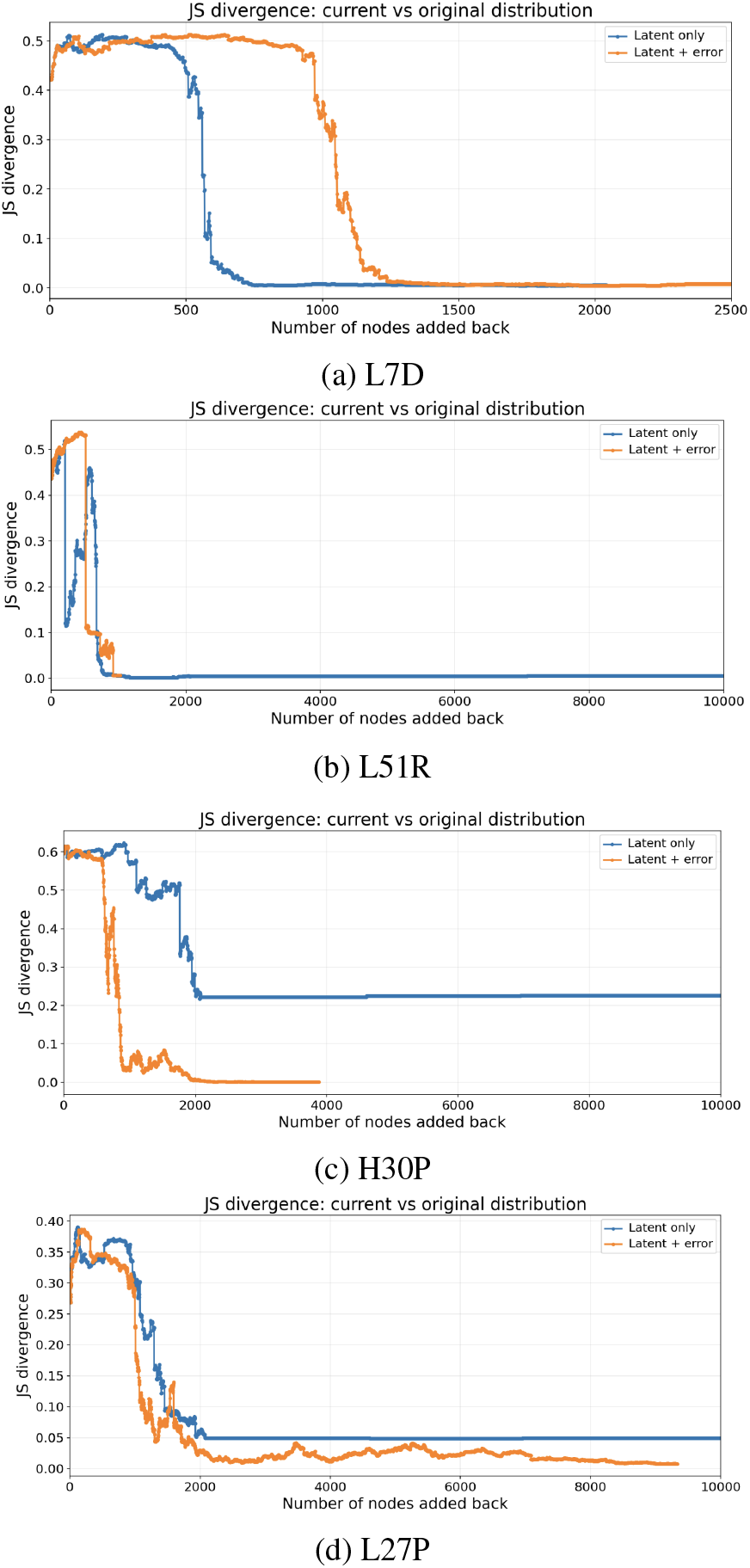
JS-divergence recovery trajectories at layer 16 for four representative mutations, plotting the Jensen–Shannon divergence between the recovered and clean output distributions as latent–token nodes are restored in descending order of integrated-gradient attribution (latent-only, blue; latent-plus-error, orange). **Horizontal axes not shared:** L7D (a) spans ~ 2,500 restored nodes, while L51R (b), H30P (c), and L27P (d) extend to 10,000 in the x-axis.

#### F.3 Distribution Snapshots

Figure 8 shows the amino-acid distribution at the masked L7 position as the recovered circuit *P*_*C*_ is built up toward the full-model distribution *P*_*M*_, for the L7D mutation at layer 16 (latent-only circuit). At a JS-recovery of ≥ 0.15 (16 nodes), *P*_*C*_ remains nearly flat across the twenty amino acids and fails to reproduce the sharp leucine peak of *P*_*M*_ (≈ 0.92), leaving a residual JS divergence of ~ 0.50. By JS-recovery ≥ 0.50 (670 nodes), *P*_*C*_ develops a clear leucine peak (≈ 0.70) but still falls below the clean peak by ~ 0.22 probability units. At JS-recovery ≥ 0.90 (821 nodes), *P*_*C*_ and *P*_*M*_ are nearly indistinguishable, where a sparse subset of 821 of the 2,080 active latents (39.5%) is able to reproduce the full output distribution at layer 16. Similar trends hold at other layers showing that the latents restored during recovery using JS-recovery score restores the most important components of the network necessary for predicting mutation-effects by restoring almost the original probability distribution at the masked position *t*.

**Figure 8:**
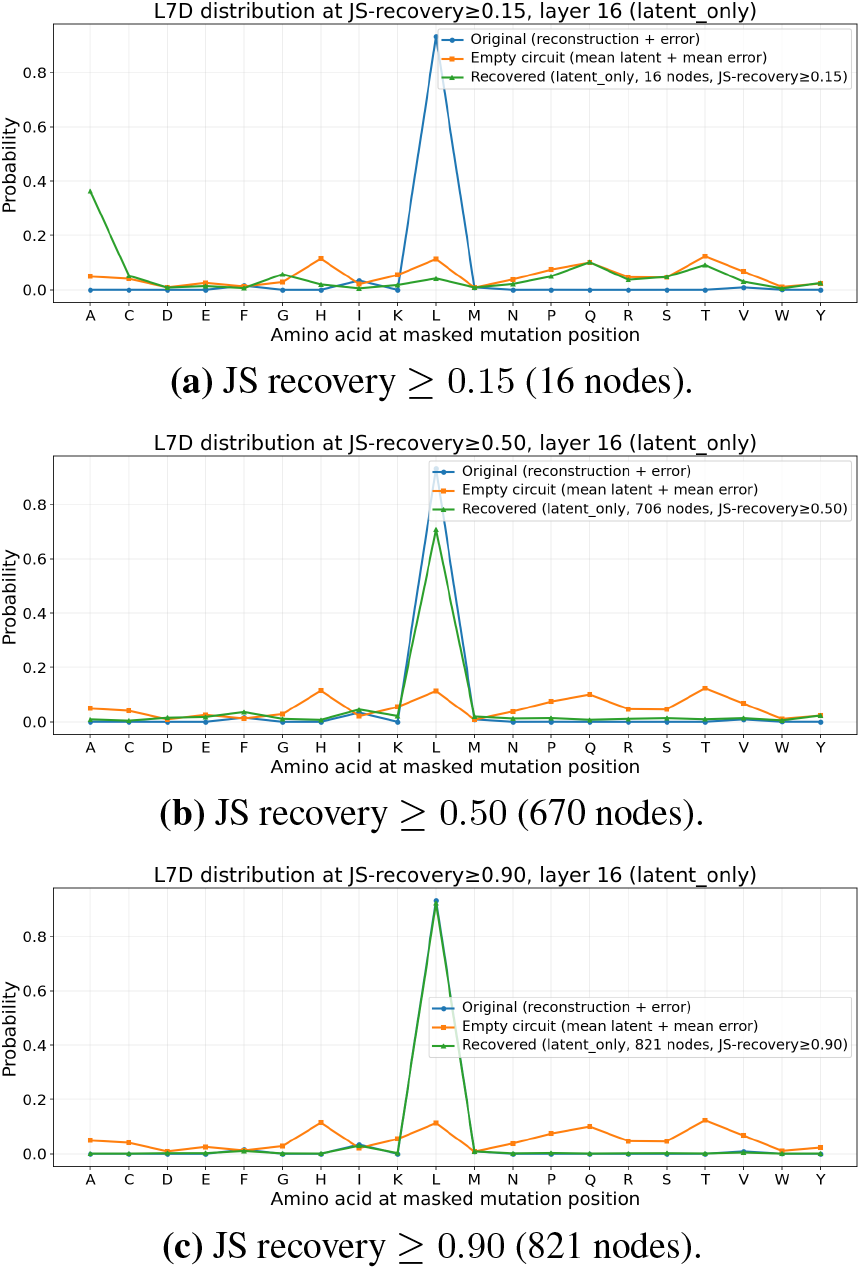
Amino-acid probability distributions at the masked mutation position for the L7D layer-16 latent-only circuit at increasing JS-recovery thresholds. The original-model distribution *P*_*M*_, recovered distribution *P*_*C*_, and empty-circuit distribution *P*_∅_ are shown in blue, green, and orange, respectively. (a) *P*_*C*_ remains diffuse and does not recover the leucine peak in *P*_*M*_ at *θ* = 0.15. (b) *P*_*C*_ develops a clear leucine peak while approaching *P*_*M*_ at *θ* = 0.50. (c) *P*_*C*_ and *P*_*M*_ are nearly indistinguishable at *θ* = 0.90.

#### G ToolUniverse-Based Latent Labeling Pipeline

We customized the ToolUniverse `ESM_describe_sae_feature` procedure to describe the layer-16 SAE latents selected by our JS–IE circuit analysis. The original evidence-aggregation procedure was retained: each latent was evaluated on a fixed ten-protein panel, its strongest activating residues were matched to UniProt annotations, and a coarse label was assigned using a dominant-category vote. The principal modification was to replace the original ESM-C SAE inference backend with our ESM-2-650M layer-16 SAE. The labeling procedure was deterministic and did not use a language model.

#### Selection of latents for description

The input to the labeling pipeline was the ranked set of latent–token nodes obtained using JS–IE. Nodes were ordered by the absolute magnitude of their indirect-effect attribution, and duplicate latent IDs occurring at different token positions were removed while preserving the rank of their first occurrence. In the reported analysis, we described the first *N* = 15 unique latents. The JS–IE scores and token positions determined only which latents were submitted to the labeling procedure; they did not contribute to the label assigned to a latent.

#### Customized SAE activation extraction

We evaluated each selected latent on the fixed ToolUniverse panel of ten proteins: TP53, EGFR, KRAS, thrombin, CYP3A4, insulin, hemoglobin beta, tissue plasminogen activator, albumin, and AKT1. For each protein, the canonical sequence was passed through ESM-2-650M, and its layer-16 residue representations were encoded using our 4096-dimensional SAE. Let *a*_p,r,j_ denote the activation of latent *j* at residue *r* in protein *p*. Following the ToolUniverse procedure, we retained the three residues with the largest absolute nonzero activation for each latent in each protein:

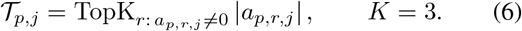

Thus, each latent contributed at most 30 residue examples across the ten-protein panel, while proteins in which the latent did not activate contributed no examples.

#### Mapping UniProt evidence to categories

For every selected residue, we retrieved all UniProt features whose annotated range overlapped that position. Informative UniProt feature types were mapped to the same coarse categories used by the original ToolUniverse protocol: `Active site` and `Site` were mapped to catalytic; `Binding site`, `Metal binding`, and `DNA binding` to ligand binding; `Modified residue`, `Cross-link`, `Glycosylation`, and `Lipidation` to post-translational modification; `Domain`, `Repeat`, and `Zinc finger` to domain; and `Motif` to motif. `Disulfide bond` and `Coiled coil` were mapped to structural stability, whereas `Helix`, `Beta strand`, and `Turn` were mapped to secondary structure. Transmembrane, signal-peptide, and propeptide annotations were retained as separate categories. General annotations such as `Region`, `Chain`, `Natural variant`, `Mutagenesis`, and `Alternative sequence` were recorded in the evidence table but excluded from category voting. The voting procedure to the mapped category remains the same as the original ToolUniverse procedure.

#### Assignment of the proposed label

The dominant category was defined as the category receiving the largest number of informative votes:

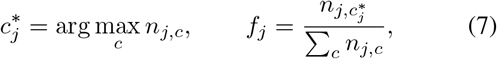

where *f*_j_ is reported as the `dominant_vote_fraction`. This quantity is a heuristic measure of agreement among the retrieved annotations and is not a calibrated confidence probability. When multiple categories received the same maximum number of votes, the output retained one dominant category while setting `dominant_category_tie` to true and recording all tied categories. Latents with no informative category votes were assigned the category `uncategorized`. The `proposed_label` column was produced by applying a fixed human-readable mapping to 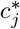. For example, the categories `secondary-structure`, `structural-stability`, `domain`, and `ligand-binding` were converted to `secondary-structure-associated feature`, `structural-stability-associated feature`, `protein-domain-associated feature`, and `ligand-binding-associated feature`, respectively. Analogous mappings were used for catalytic sites, post-translational modifications, motifs, transmembrane regions, signal peptides, and propeptides. Formally,

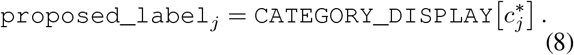

Alongside each label, we report the number of panel proteins in which the latent activated, the number of selected residue examples overlapping at least one informative annotation, the number lacking informative annotations, the complete category vote counts, the dominant-vote fraction, and any dominant-category tie. We also retained the exact residues, activation magnitudes, UniProt feature descriptions, and annotation ranges used to produce each vote, enabling the aggregate labels to be audited.

### H Circuit Interpretation

#### H.1 Top Five Latents by Method

Table 1 lists the five highest-ranked layer-16 latents under indirect-effect (IE) and activation-difference (AD) ranking for each mutation. For the buried mutations L7D and L51R, IE preferentially ranks latents associated with residues in 3D contact with the mutation site, together with several unresolved or diffuse features. In contrast, AD primarily selects residue-identity features corresponding to the wild-type or mutant amino acid, such as leucine, aspartate, and arginine detectors, along with several latents lacking a clear interpretation. For H30P, the IE-ranked contact-associated latents highlight residues near L7 and L51 rather than H30, indicating that the recovered circuit is less specifically aligned with the local environment of this mutation. Overall, IE more consistently identifies structurally relevant features for the buried mutations, whereas AD is dominated by activation changes associated with residue identity.

**Table 1:** Five highest-ranked layer-16 latents under indirecteffect (IE) and activation-difference (AD) ranking for each mutation.

| Mutation | Method | Rank | ID | Proposed label |
| --- | --- | --- | --- | --- |
| L7D | IE | 1 | 1695 | Contact with mutation site |
|  | IE | 2 | 3454 | Unresolved/diffuse |
|  | IE | 3 | 888 | Contact with mutation site |
|  | IE | 4 | 1075 | Unresolved/diffuse |
|  | IE | 5 | 3509 | Contact with mutation site |
|  | AD | 1 | 2339 | Leucine detector highlighting the mutation site |
|  | AD | 2 | 2329 | Aspartate detector highlighting the mutant residue |
|  | AD | 3 | 454 | No clear interpretation |
|  | AD | 4 | 633 | No clear interpretation |
|  | AD | 5 | 1001 | Contact with mutation site |
| L51R | IE | 1 | 888 | Contact with mutation site |
|  | IE | 2 | 1075 | Unresolved/diffuse |
|  | IE | 3 | 1399 | Contact with mutation site |
|  | IE | 4 | 1420 | Contact with mutation site |
|  | IE | 5 | 1370 | Unresolved/diffuse |
|  | AD | 1 | 3953 | Arginine detector |
|  | AD | 2 | 2339 | Leucine detector |
|  | AD | 3 | 1237 | Leucine detector |
|  | AD | 4 | 2024 | No clear interpretation |
|  | AD | 5 | 813 | No clear interpretation |
| H30P | IE | 1 | 1695 | Contacts L7 and L51, but not H30 |
|  | IE | 2 | 3454 | Unresolved/diffuse |
|  | IE | 3 | 3509 | Contacts L7 and L51, but not H30 |
|  | IE | 4 | 1075 | Unresolved/diffuse |
|  | IE | 5 | 888 | Contacts L7 and L51, but not H30 |
|  | AD | 1 | 3713 | No clear interpretation |
|  | AD | 2 | 1160 | Histidine detector |
|  | AD | 3 | 3426 | No clear interpretation |
|  | AD | 4 | 2628 | No clear interpretation |
|  | AD | 5 | 1888 | No clear interpretation |

#### H.2 Divergent Concept Latents

Table 2 displays some latents whose global InterProt and Tool-Universe descriptions diverge from the role the latent plays in the DNAJA1 mutation-effect circuit. The “proposed label” column reports our structure-guided, mutation-specific interpretation, assigned by manual inspection of each latent’s activating residue positions relative to the mutation site in the DNAJA1 structure according to the categories described in (Appendix H.3). These cases illustrate a latent’s corpus-derived interpretation does not always predict which residues carry a causal signal for a specific prediction. L27P does not have any mutation crossing the recovery threshold, hence is eliminated from the analysis.

**Table 2:** Latents whose global InterProt descriptions differ from their mutation-specific roles in the DNAJA1 circuit.

| Mutation | Latent | InterProt description | DNAJA1 circuit annotation | ToolUniverse label |
| --- | --- | --- | --- | --- |
| L7D | 1695 | Strongly activates in SAM domains | Contact with the mutation site at Y22 and Y48 | Secondary-structure-associated feature |
| L7D | 955 | Top Pfam associations are kinase-related: PF00069 (protein kinase), PF07714 (tyrosine and serine/threonine kinase), and PF03109 (ABC1 kinase-like) | Contact with the mutation site at Y22 | Secondary-structure-associated feature |
| L7D | 2695 | Top Pfam associations correspond to ATPase and transporter families: PF00005 (ABC transporter), PF13476 (AAA domain), and PF02463 (RecF/RecN/SMC N-terminal domain) | Contact with the mutation site at Y22 and Y48 | Secondary-structure-associated feature |
| L7D | 3810 | Top Pfam association is PF00069 (protein kinase domain) | Contact with the mutation site at Y48 | Structural-stability-associated feature |
| L7D | 2187 | Top-activating sequences are paraneoplastic or paraneoplastic-like antigens | Contact with the mutation site at Y22 | Secondary-structure-associated feature |

#### H.3 Annotation Categories and Assignment Criteria

We assigned each latent to one mutually exclusive annotation category using the spatial distribution of its absolute integrated-gradients attribution across residue positions. For latent *j* in mutation context *m*, let 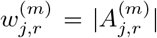 denote its attribution at residue *r*. Each residue position was first assigned to exactly one of the following categories:

- **Mutation site:** the masked mutation position *t*.
- **3D contacts:** residues included in the precomputed structural-contact set for position *t*, excluding the mutation site itself. We retained contacts with a contact fraction of at least 0.8.
- **Sequence-local:** residues within ±5 sequence positions of *t* that were not assigned to the mutation-site or 3D-contact categories.
- **HPD/motif:** residues at positions 30–32, corresponding to the conserved HPD motif, that were not assigned to a higher-priority category.
- **Other residues:** all remaining residue positions.

Because these annotations can overlap, positions were assigned using the following fixed priority:

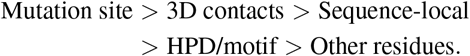

Thus, each attributed residue contributed to exactly one category. The attribution assigned to category *c* for latent *j* was

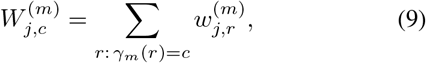

where *γ*_*M*_(*r*) is the exclusive category assigned to residue *r*. We normalized these weights within each latent to obtain its category profile,

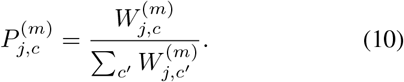

For a mutation-specific latent, the assigned annotation was the category with the largest normalized attribution in its corresponding mutation context. For a latent shared between mutations *m*_1_ and *m*_2_, we first averaged its normalized category profiles across the two contexts,

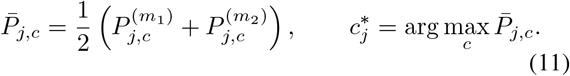

This gives equal weight to the two mutation contexts rather than allowing the context with greater total attribution magnitude to dominate the label. Contexts with no usable attribution for a latent were omitted from the average. Exact ties were resolved using the category priority given above.

We additionally defined an **Unresolved/diffuse** category for latents whose attribution did not concentrate on a coherent spatial or residue-identity pattern. A latent was assigned to this category when it activated at five or more residue positions, none of the mutation-site, 3D-contact, sequence-local, or HPD/motif categories accounted for at least 75% of its total absolute attribution, and no single amino-acid identity accounted for at least 75% of that attribution. The residual *Other residues* category was excluded from this diffused category calculation because it combines heterogeneous positions rather than representing a single biological concept. For a shared latent, the unresolved/diffuse label was applied if these criteria were satisfied in either mutation context. This label overrides the category selected by the maximum-attribution rule.

Finally, each latent’s complete absolute-attribution mass was assigned to its single selected category. Consequently, the reported category fractions quantify the distribution of attribution across exclusively labeled latents and do not double-count latents or residue-level attribution across overlapping annotations.

### I Shared Circuit Reuse and Enrichment

#### I.1 Within-Family Jaccard Distributions

Table 3 shows the median within-family Jaccard similarity (%) of recovered layer-16 circuits at the *τ* = 0.8 recovery threshold, across ESM-2 layers for Figure 4(a).

**Table 3:** Median within-family Jaccard similarity (%) between recovered mutation circuits at the *τ* = 0.8 recovery threshold. Each entry is the median across resolvable mutation pairs within the corresponding family and ESM-2 layer. Dashes indicate layers for which the required none of the constitutent mutations reached the threshold.

| Mutation family | ESM-2 layer |  |  |  |  |  |  |  |
| --- | --- | --- | --- | --- | --- | --- | --- | --- |
|  | 4 | 8 | 12 | 16 | 20 | 24 | 28 | 32 |
| H30P-H30X | – | 50.7 | 47.7 | 100.0 | 47.3 | 49.5 | 53.7 | 43.7 |
| L27P-L27X | 75.3 | – | 40.0 | 100.0 | 64.4 | 62.0 | 100.0 | 2.8 |
| L51R-L51X | 71.9 | 59.4 | 38.3 | 68.3 | 58.5 | 46.2 | 40.0 | – |
| L7D-L7X | 62.7 | 51.5 | – | 64.0 | 55.0 | 55.5 | 45.1 | 39.6 |

#### I.2 Enrichment Calculations for Biological Annotations

We quantified whether each biological annotation was preferentially represented in latents shared between two mutation circuits relative to latents specific to either mutation. Enrichment was computed independently for every mutation pair and SAE layer using the mutually exclusive latent annotations defined in Section H.3. Let *S* denote the shared latents for mutations *m*_1_ and *m*_2_, and let 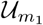 and 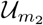 denote the latents specific to each mutation.

For latent *j* in mutation context *m*, we first computed its total absolute attribution across residue positions,

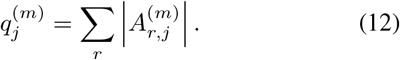

Because each latent was assigned to exactly one category, its complete attribution mass 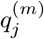 contributed to that category. For a latent set *G*, the attribution fraction assigned to category *c* in context *m* was

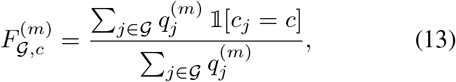

where *c*_j_ is the assigned category of latent *j*. Shared latents used the same pair-level category in both mutation contexts, whereas mutation-specific latents were labeled using their corresponding mutation context.

To prevent one mutation from dominating because of differences in total attribution magnitude, we averaged the normalized fractions from the two mutation contexts. The shared-circuit fraction was

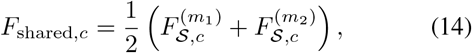

and the mutation-specific reference fraction was obtained by averaging 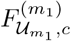 and 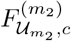. When one mutation had no specific latents, only the available mutation-specific context was used. Annotation enrichment was then defined as

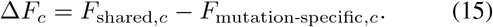

Thus, Δ*F*_*C*_ *>* 0 indicates that category *c* accounts for a larger fraction of attribution in the shared circuit, whereas Δ*F*_*C*_ *<* 0 indicates relative depletion.

We estimated uncertainty using a latent-level bootstrap with 200 replicates. Within each shared or mutation-specific latent set, latents were sampled with replacement while preserving the original set size, and the annotation fractions and Δ*F*_*C*_ were recomputed. The 2.5th and 97.5th percentiles of the resulting distribution defined the 95% confidence interval. An enrichment value was marked as distinguishable from zero when this interval lay entirely above or below zero. Pair–layer cases lacking the required recovered-circuit or attribution outputs, shared latents, or any mutation-specific comparison were treated as unavailable.

For mutation-family summaries, enrichment was first calculated separately for each evaluable mutation pair. We then reported the median Δ*F*_*C*_ across pairs within a family and layer, with the interquartile range describing pair-level variability. Latents were therefore never pooled across different mutation pairs before computing enrichment.

#### I.3 Other Biological Annotation Enrichment Plots

Figure 9 displays the enrichment plots for other biological annotations. Enrichment plots for 3D contacts and Unresolved/diffused category are displayed in Figure 4. Unlike layer 16 showed contact enrichment for specific mutation families, for all of these remaining annotations there’s no such high enrichment for any of the annotations. Interestingly, in layer 12 L27X family shows enrichment for the “Other residues” category, however, on closer inspection, those latents didn’t show any clear interpretations.

**Figure 9:**
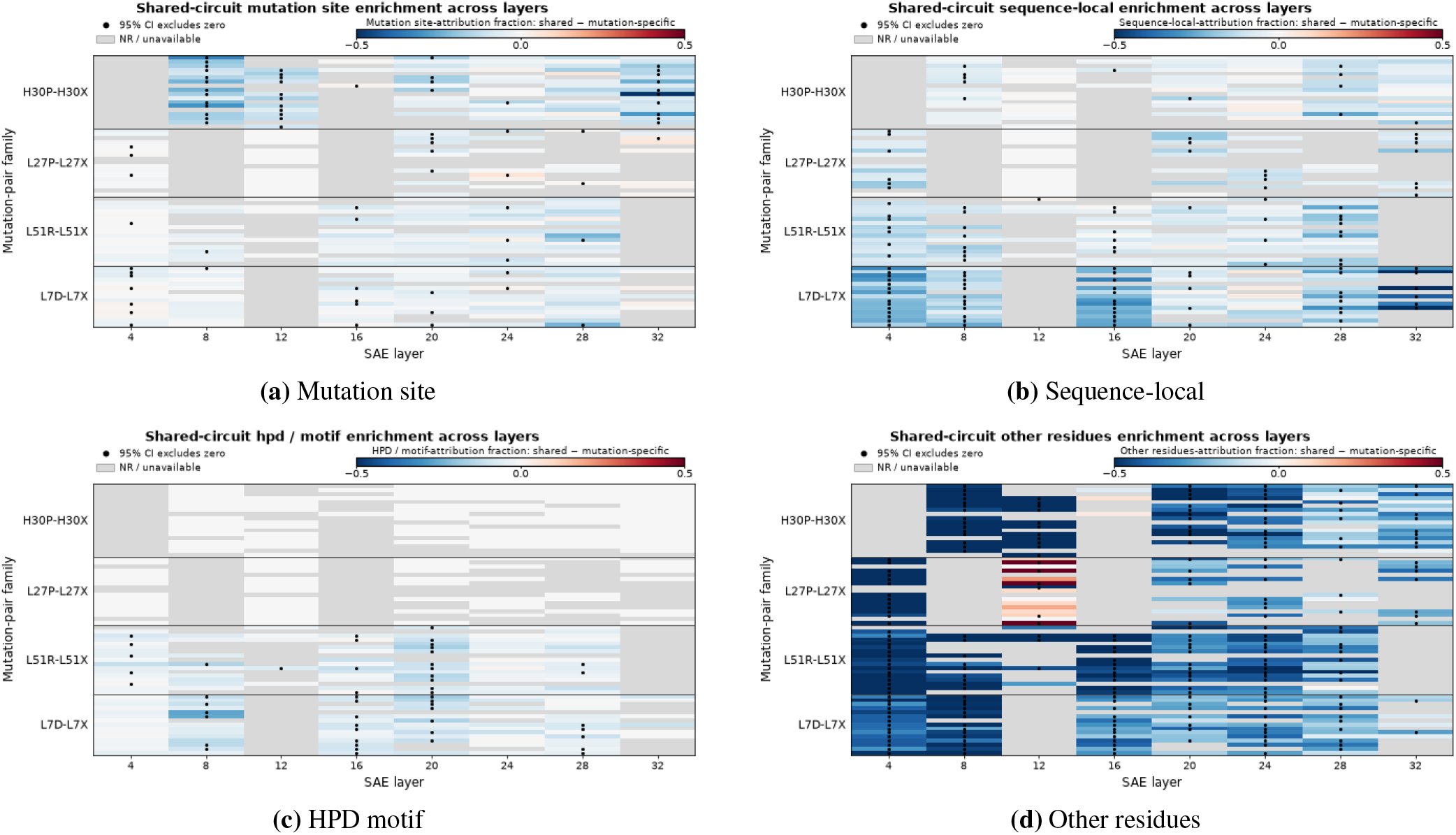
Circuits associated with different biological annotation categories. (a) Mutation-site latents. (b) Sequence-local latents. (c) HPD-motif latents. (d) Latents associated with other residues.

